# Experimental manipulation of body size alters senescence in hydra

**DOI:** 10.1101/2020.06.19.161034

**Authors:** Kha Sach Ngo, Berta Almási, Zoltán Barta, Jácint Tökölyi

## Abstract

Body size has a fundamental impact on the ecology and physiology of animals. Large size, for instance, is often associated with increased fecundity and reproductive success. A persistent correlation between body size and individual longevity is also observed across the animal world, although this relationship proved difficult to understand due to the inseparability of body size from growth rate and the widespread collinear relationship between body size with other life history traits. Here, we used *Hydra oligactis*, a freshwater cnidarian with high tissue plasticity and inducible ageing as an experimental system to understand the causal roles of body size on reproduction and senescence. We first show that large size predicts accelerated sexual development, increased fecundity and reduced survival in a population sample of this species kept under common garden conditions in the laboratory. Next, using phenotypic engineering, we experimentally increased or decreased body size by reciprocally grafting pieces of the body column differing in size between hydra polyps. Experimentally reduced body size was associated with delayed sexual development and reduced fecundity. In parallel, post–reproductive survival was significantly higher in polyps with reduced size. These results suggest that small hydra can physiologically detect their reduced body size and adjust reproductive decisions to achieve a higher post–reproductive survival. Our observations offer a new perspective on why smaller individuals within a species live longer by suggesting a growth–independent link between body size, reproduction and senescence.

## INTRODUCTION

Does body size *per se* control critical aspects of life history? Allometry, the scaling of biological patterns and processes to body size, has fascinated scientists for centuries. Body size influences most aspects of biology including physiological performance, predator–prey interactions, life-history characters, ecological patterns, and evolutionary trajectories (Peters 1983, Damuth 2007). A larger body size provides competitive advantage (Darwin 1874, Arak 1988, Andersson 1994, Székely et al. 2000), and consistently correlates with a disproportionately higher reproductive output (Peters 1983, Allaine et al. 1987, Reiss 1987, Hendriks and Mulder 2008, Barneche et al. 2018).

In addition to fecundity, a persistent relationship between body size and individual longevity has also been observed (reviewed in Austad 2010), although the underlying mechanisms are complex and often hotly debated. On the one hand, a large body size implies better condition, increased resources against starvation and more protection from predators, resulting in some cases in a positive relationship between individual size and longevity. For instance, a positive relationship between body size and longevity has been described in field populations under resource limitation (e.g. Forsman, 1993, Gaillard et al. 2000) and in comparisons between body size and longevity across species (e.g. Promislow & Harvey 1990, Speakman 2005). On the other hand, larger individuals bear the viability costs of longer development and increased growth rates and the metabolic costs of maintaining a larger body (Blanckenhorn 2000, Metcalf and Monaghan 2003). Therefore, when variation in resource availability is reduced (such as in a laboratory environment), the relationship between body size and longevity within species is generally a negative one: smaller individuals tend to live longer (Austad 2010). In laboratory rats and mice, for instance, body size correlates negatively with longevity across strains (Rollo 2002). A similar correlation is seen across dog and horse breeds (Austad 2010, Kraus et al. 2013, Bartke 2017). Reduction or deficiency in growth hormone (GH) or Insulin–like Growth Factor–1 (IGF–1) and experimental dietary restriction (DR) in mice have produced smaller animals with substantially longer lifespan than co–specifics from the same strain (Bartke 2017). Even in humans, short stature has been shown to predict increased longevity in some analyzes (Samaras et al. 2003, He et al. 2014), although this relationship is complicated by the contrasting effects of height on different classes of diseases (Austad 2010).

As the previous findings show, body size is an important predictor of fecundity and longevity/ageing. Despite the importance of these relationships, most previous studies relied, unsurprisingly, on correlations between body size and life history traits. Previous attempts to experimentally manipulate body size, such as DR, GH manipulation, or genetic mutants have not provided sufficient evidence whether it is body size itself that impacts reproduction or longevity, as all of these methods influence body size through modifying growth rate. Accelerated growth is known to have negative effects on survival (Metcalf & Monaghan 2003) and could be the mechanism behind the body size – longevity relationship. However, it is unclear whether body size can influence senescence through growth–independent mechanisms. Moreover, engineering transgenic strains, e.g. in the insulin/IGF–1 signaling pathway, can introduce unintentional side effects as insulin/IGF–1 signaling plays a key role in controlling many physiological processes other than size and aging, including protein synthesis and glucose metabolism (e.g. McCulloch & Gems 2003). These factors combined complicate conclusions that draw a causal effect between body size and ageing using data obtained from genetic mutants and DR–strains. Therefore, it is necessary to examine the effect of direct experimental body size manipulation, isolated from genetic and environmental factors, to determine if body size itself determines fecundity and ageing.

Here, we present our results obtained from direct experimental manipulation of body size on an emerging model system in ageing research, the freshwater cnidarian *Hydra oligactis* (Pallas 1766). This system is an ideal candidate for this experiment, due to several characteristics. Hydra as a genus is renowned for their regenerative capability, providing a reliable platform for surgical manipulation to change body size with negligible risk in causing debilitating injuries or death. *H. oligactis* in particular exhibits unique sexual behaviours different from other well–established Hydra species, wherein sexual reproduction results in catastrophic senescence and higher mortality rates compared to asexually reproducing strains (Martínez 1998, Yoshida et al. 2006, Schaible et al. 2015, Tökölyi, et al. 2017, Sebestyén et al. 2018, Tomczyk et al. 2020). Using this system we first show that body size predicts sexual development and post–reproductive survival rate in a population sample maintained under standard conditions in the lab. Next, we conducted size manipulation on temperature–, age– and diet–standardized presexual *Hydra oligactis* to produce 3 distinct treatment groups, enlarged, control and reduced. Following the induction of sexual development, the polyps were maintained and monitored at regular intervals for quantities of gonads (representing sexual development) and survival, respectively.

## MATERIALS AND METHODS

### Field collection of Hydra strains and common garden experiment

Hydra strains, the population sample hereafter, were collected from an oxbow lake near Tiszadorogma, Hungary (47.67 N, 20.86 E). Sampling was performed 4 times in two subsequent years (31th May 2018, 1st Oct. 2018, 16th May 2019, 24th Sept. 2019). On each collection date hydra polyps were collected from multiple locations within the lake, brought to the laboratory and propagated asexually for 10 weeks while keeping them individually under standard conditions (constant temperature of 18 ºC, 12/12 h dark/light cycle, artificial hydra medium, feeding 2x/week; see Tökölyi et al. 2020 for a more detailed description). After 10 weeks, temperature was lowered to 8 ºC and photoperiod changed to 8/16 h dark/light cycle to stimulate autumn conditions and induce sex. The presence and number of gonads and survival was recorded throughout this period for five months and polyps were fed and cleaned 2x/week.

For the population sample, we examined altogether N = 1584 specimens. A subset of these individuals (N = 330) died during the asexual propagation phase or after cooling, before producing either buds or gonads and hence generating usable data. We further excluded N = 18 individuals belonging to 2 strains that contained sex–changed animals. Final sample size was therefore N = 1236 polyps belonging to N = 181 strains. However, there was missing data in some of the variables measured: 1) we accidentally lost N = 12 post–reproductive individuals and could not quantify their survival and 2) we failed to measure body size in N = 16 cases (e.g. because the polyp was highly contracted). Therefore, sample sizes in some analyses are slightly lower.

### *H. oligactis* strains used for size manipulation

We used one strain of male (C2/7) and one strain of female (X11/14) *H. oligactis* for experimental manipulations (see Sebestyén et al. *in press*, for a detailed description of these strains). Both strains derive from two polyps that were collected from the same population earlier (September 2016) and were maintained asexually in the lab since then. The polyps were housed in 6–well tissue culture plates in a climate chamber at 18 °C and 12/12 hours light–dark cycle.

The buds were propagated to create a constantly renewing population of about 60 for each strain. Each polyp was housed and identified in individual wells, its age and strain identification number recorded. Maintenance was carried out 4 times per week (on Mondays, Tuesdays, Thursdays, Fridays), during which the adult polyps were fed and cleaned. We maintained the strains in an age–standardized manner: polyps reaching 3 weeks of age were removed from culturing and exchanged with one of their detached buds. The adults were discarded while the buds remained in the strain population. This regular replacement was maintained for 8 months prior to the experiment, hence experimental animals derive from an asexual lineage where the asexual parent, grandparent, great–grandparent, etc. were approximately of the same age (3 weeks). After 8 months the adult polyps removed from rearing were retained for experimental treatment.

### Experimental size manipulation

Polyps that were 3–4 weeks old and derived from the age–standardized strains were randomized into 4 experimental treatment groups per set, enlarged, reduced, control 1 and control 2. Polyps with obvious abnormalities and/or signs of stress, were discarded. Polyps were photographed prior to and after experimental treatment using a Euromex StereoBlue microscope, a Euromex camera with a standard 1–mm grid sheet beneath the polyp. Pre–treatment and post–treatment body surface area of each polyp was measured as a proxy for body size, using the software ImageJ (Schneider et al. 2012).

Polyps of each treatment group received 2 transverse cuts using a sharp medical scalpel to create 3 distinct parts containing the head, mid–body ring, and lower–body plus foot peduncle. The two control polyps were cut so that their mid–body rings were of similar sizes, and the rings were then exchanged between the two polyps. The enlarged and reduced polyps underwent similar procedures, except for the body parts being cut at different lengths so that the enlarged polyp would exchange its small mid–body ring for the much larger body ring from the reduced polyp (and vice versa for the reduced). After the exchange of rings, the 3 parts of each polyp, the host’s head, donor’s ring, and host’s foot peduncle, were secured together with a small glass capillary needle to let heal the cuts for 2–3 hours on room temperature. The polyps were then photographed for post–treatment body size measurements 24 h after healing and moved to a climate chamber with 8 °C and a 16/8 hours dark/light cycle to induce sex. They were fed and cleaned twice per week during the cold phase.

All grafting was done among individuals belonging to the same strain (i.e. we didn’t graft tissue from the male to the female strain or vice versa). Out of 59 sets of 4 polyps (n = 236 individuals in total), we excluded pairs that contained polyps of failed experimental attempts and those that exhibited abnormal sexual behaviors, i.e. sex change or remained asexual, were excluded from the data. Final sample size was N = 90 male polyps (N = 24 reduced, N = 42 control and N = 24 enlarged) and N = 92 female polyps (N = 22 reduced, N = 50 control and N = 20 enlarged).

### Quantifying appearance and number of gonads

Following cooling, animals were monitored for the appearance of gonads during routine maintenance. In males, testes develop simultaneously around the whole body column, but we counted mature testes only on one side of the body column, without rotating the polyp (to avoid counting gonads twice). For each individual we used the maximum number of testes observed during the whole reproductive period as a proxy for male fecundity. In females, eggs develop sequentially and unfertilized eggs detach from the parent. We counted and removed detached eggs during routine maintenance and used the sum of detached eggs as a proxy for female fecundity. In a few cases, we noticed mature eggs on the body column of polyps in the populations sample but didn’t find them detached in the tissue culture plates, likely because unfertilized eggs can sometimes degrade. These eggs were not included in our estimation of female fecundity but we still scored the polyps involved as females.

To gain insight into size–specific reproductive allocation, we also calculated relative gonad number by dividing the no. gonads with body size for both males and females.

### Quantifying survival

In *H. oligactis* sexual reproduction is followed by post–reproductive senescence during which movement and feeding ability declines, polyps become unresponsive to touch, their body becomes discolored and shrinks in size, ultimately resulting in an amorphic, necrotic mass. In the lab strains originally characterized in detail by Yoshida et al. (2006), all individuals show these symptoms within three months. In another lab strain (Ho_CR) characterized by Tomczyk et al. (2020), polyps develop sexually but do not show morphological signs of senescence. The hydra strains derived from our source population show an intermediate pattern, wherein most polyps undergo post–reproductive senescence, but some of them are able to regenerate from this senescent state and resume asexual reproduction (Sebestyén et al. i*n press*). This reversal of senescence probably rests on the ability of hydra polyps to regenerate whole bodies from just a few cells (Bosch 2007).

To avoid scoring animals dead when they retain the ability to survive, we kept our experimental animals for longer (5 months in the common garden study, 6 months in the size–manipulation experiment), even if they shrank to a very small size and looked a mass of amorphic necrotic tissue. Animals were scored “survived” if they regenerated and produced asexual buds following sexual reproduction in both experiments. Furthermore, in the common garden experiment we also scored animals “survived” if they didn’t produce buds but had intact tentacles, looked healthy and were able to feed at the end of the five months cooling period. No such individual was observed in the size–manipulation experiment (i.e. they either produced buds, disappeared or remained senescent by the end of the experiment). Animals that shrinked in size and disappeared or lost their normal body shape and became an amorphic mass of necrotic tissue were scored as “not survived”.

### Statistical analyses

In the population–level study we used Linear Mixed Effects Models (LMM) or Generalized Linear Mixed Models (GLMM) to analyze the effect of body size on: 1) the probability of sexual reproduction (vs. remaining asexual during the whole cooling phase; binomial GLMM), 2) start of sexual reproduction (LMM), 3) maximum number of testes per male and total number of eggs per female (Poisson GLMM), 4) size–standardized gonad number (LMM) and 5) survival (combined for sexuals and asexuals and separately for males and females; binomial GLMM). We log–transformed time to gonadogenesis to improve normality. Strain ID was included as random effect and we controlled for polyp age and collection date in all models. Body size and polyp age were scaled to zero mean and unit variance to improve model convergence. An observation–level random effect was added to the model analyzing the no. eggs in females to control for overdispersion of this variable.

We used LMM or GLMM to analyze the effect of experimental treatment on: 1) post–treatment polyp size, 2) start of sexual reproduction, 3) maximum number of testes per male and total number of eggs per female, 4) size–standardized gonad number and 5) survival. The two control groups were pooled for this analyzis. Size, time to gonadogenesis and size–standardized gonad numbers were analyzed with LMM with Gaussian errors, fecundity was analyzed with GLMM with Poisson errors, while survival was analyzed with GLMM with binomial errors. We included pair ID as random effect. All analyses were done in R (v 3.6.3; R Core Team 2020), using the *nlme* package for LMM (v. 3.1–144; Pinheiro et al. 2020) and *lme4* package for GLMM (v. 1.1– 21; Bates et al. 2015). We first tested whether experimental groups are significantly different from each other via Likelihood Ratio (LR) tests. When LR tests indicated significant differences, we used the *multcomp* package in R (v. 1.4–13; Hothorn et al. 2008) to perform post–hoc comparisons with Benjamini–Hochberg correction among the experimental groups.

## RESULTS

### Body size predicts reproductive mode, timing of sexual reproduction, fecundity and post– reproductive survival at the population–level

Out of the N = 1236 individuals in the population sample 18.1% (N = 224) remained asexual throughout the 5 months of the cold phase and produced only buds. N = 393 polyps (31.8%) produced eggs, while the remaining N = 619 individuals (50.1%) produced testes. We found a significant, positive relationship between body size and the probability of sexual reproduction in response to lowering the temperature: larger individual were more likely to reproduce sexually (binomial GLMM, beta = 2.10, SE = 0.20, p < 0.001, N = 1220). There was a significant, negative relationship between body size and time to gonadogenesis, hence in larger individuals sexual development was faster (LMM, males: beta = –0.10, SE = 0.10, p < 0.001, N = 610; females: beta = –0.08, SE = 0.10, p < 0.001, N = 390; Fig. 1a, b). The number of testes or eggs produced was positively related to body size (Poisson GLMM, males: beta = 0.18, SE = 0.02, p < 0.001, N = 610; females: beta = 0.21, SE = 0.04, p < 0.001, N = 390; Fig. 1c, d). However, the size–standardized number of gonads decreased with body size (LMM, males: beta = –0.17, SE = 0.02, p < 0.001; females: beta = –0.10, SE = 0.04, p = 0.019).

**Fig. 1.**
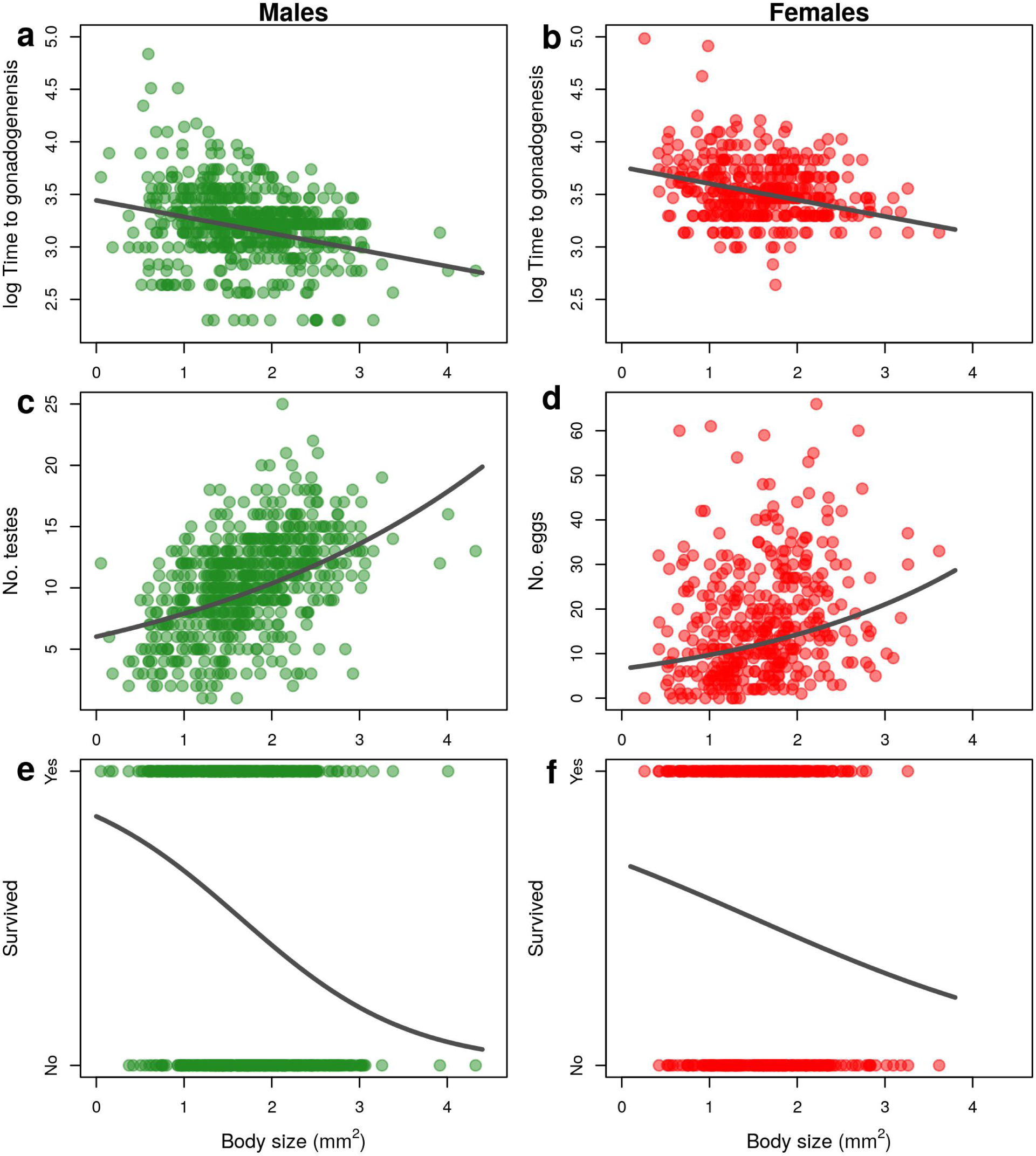
Relationship between body size and (a) start of gonadogenesis after cooling in males, (b) start of gonadogenesis after cooling in females, (c) number of testes, (d) number of eggs, (e) male survival rate and (f) female survival rate. Data points are from strains derived from populations samples of *H. oligactis* kept under common garden conditions in the laboratory. Prediction lines derive from Linear Mixed–Effects Models (a, b), Generalized Linear Mixed– effects Models with Poisson distribution (c, d) or Generalized Linear Mixed–effects Models with Binomial distribution (e, f) and contain strain ID as random factor. Sample sizes are N=610 and N=390 for male and female reproductive traits, respectively and N=604 and N=389 for male and female survival, respectively.

Overall, including both sexual and asexual individuals there was a significant negative connection between the probability of survival and body size (binomial GLMM, beta = –0.68, SE = 0.09, p < 0.001, N = 1213). The relationship was significant separately for males (binomial GLMM, beta = –0.59, SE = 0.13, p < 0.001, N = 604) and females as well (binomial GLMM, beta = –0.28, SE = 0.13, p = 0.030, N = 389; Fig. 1e, f).

### Size–manipulation

Pre–treatment body size measurements showed that the polyps we used did not significantly vary in body size prior to experimental body size manipulation (LMM, male strain, LR = 0.69, p = 0.71; female strain, LR = 1.83, p = 0.40).

In contrast, post–treatment measurements showed that our method produced polyps with significant body size differences in accordance with our 3 intended experimental groups (Fig. 2. a, b; Fig. 3.). Enlarged and reduced polyps showed larger and smaller body sizes respectively, relative to control groups’ body size (LMM, male strain, LR = 140.71, p < 0.001; control vs. reduced: beta = 0.89, SE = 0.09, p < 0.001; enlarged vs. control: beta = 0.88, SE = 0.09, p < 0.001; enlarged vs. reduced: beta = 1.76, SE = 0.10, p < 0.001; female strain, LR = 132.17, p < 0.001; control vs. reduced: beta = 0.81, SE = 0.08, p < 0.001; enlarged vs. control: beta = 0.90, SE = 0.09, p < 0.001; enlarged vs. reduced: beta = 1.71, SE = 0.10, p < 0.001).

**Fig. 2.**
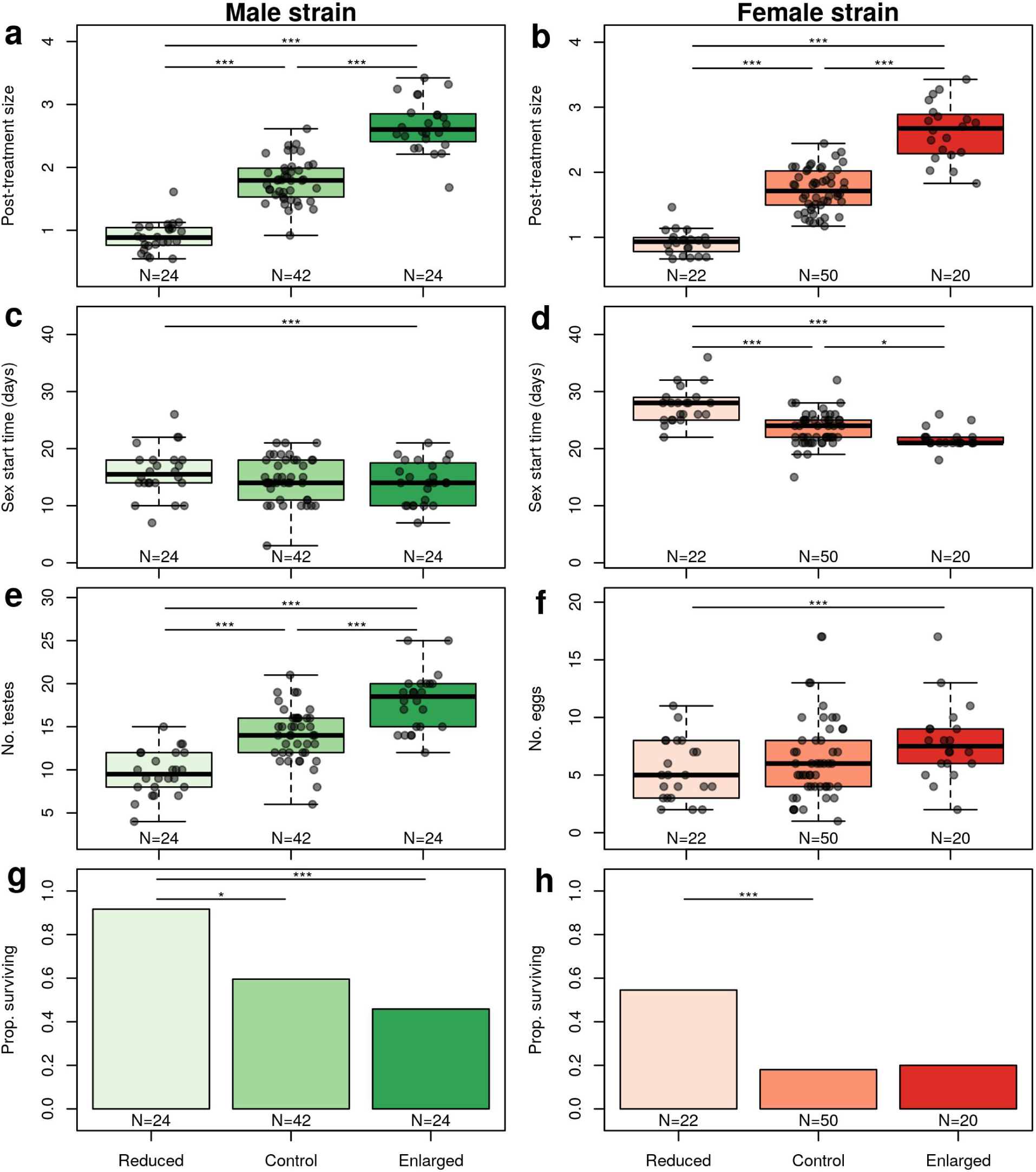
Comparison of enlarged, control and reduced individuals in post–manipulation size (body surface area measured after experimental treatment and healing; mm^2^) in the male (a) and female strain (b), start of gonadogenesis after cooling (the length of time it takes for the polyps to develop their first gonads from the beginning of cooling) (days) in the male (c) and female strain (d), no. testes in the male strain (e), no. of detached eggs in the female strain (f) survival rate (proportion of animals scored as survived after post–reproductive senescence in the male (g) and female strain (h). Comparisons between groups were done through *post–hoc* tests based on Linear Mixed–Effects Models (a–d), Generalized Linear Mixed–Effects Models with Poisson distribution (e, f) or Generalized Linear Mixed–Effects Models with Binomial distribution. Bars above pairs denote significant differences (*: p < 0.05, **: p < 0.01, ***: p < 0.001) after Benjamini–Hochberg correction. The box–and–whiskers plot shows median values, lower and higher quantiles and min–max values. Dots denote individual data points.

**Fig. 3.**
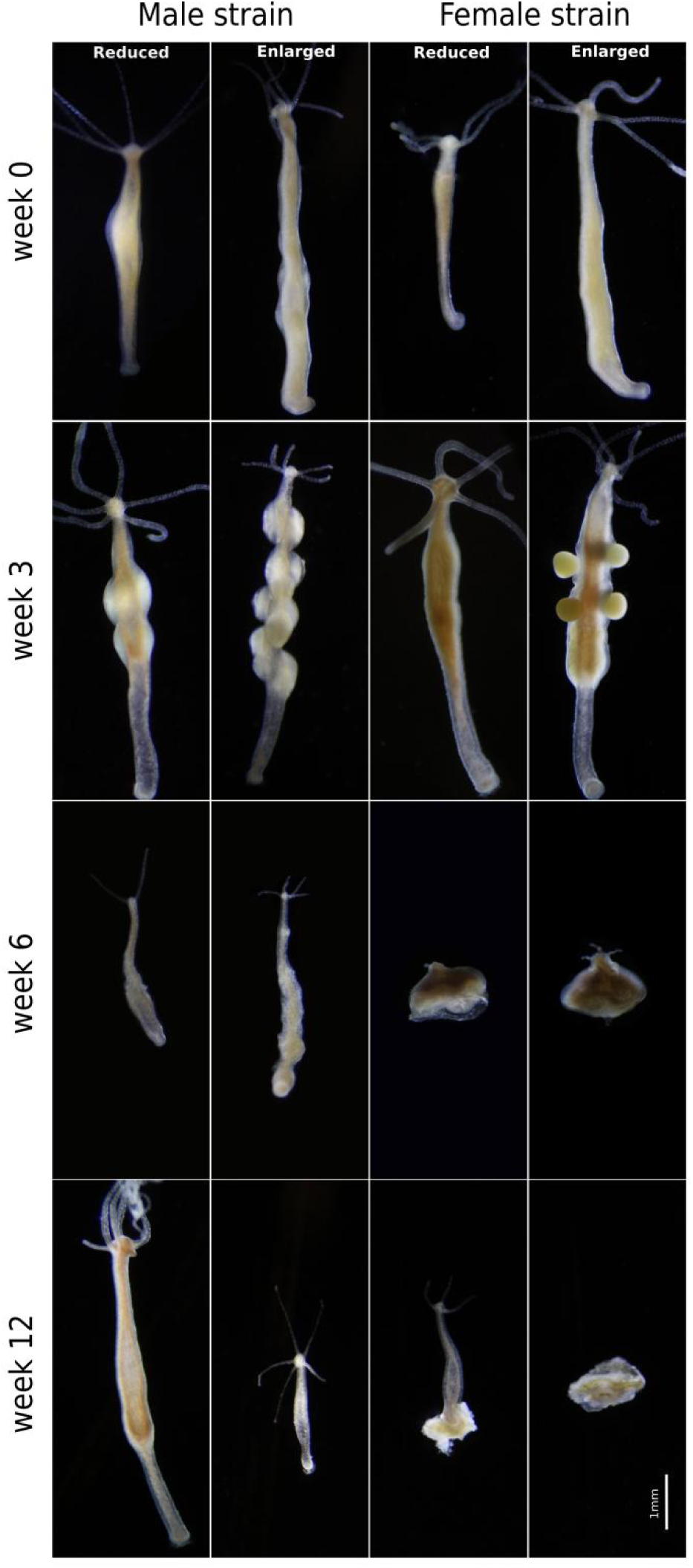
Photographs illustrating sexual development and post–reproductive senescence of reduced and enlarged males and females. Gonads are visible by week 3 except in females with reduced body size that initiate gonadogenesis latest. By week 6 all polyps show signs of post– reproductive senescence (reduced body size and shortened tentacles). At week 12, some individuals show signs of recovery. Reduced individuals are in a more advanced stage of regeneration and males are more advanced than females.

### Experimental manipulation of body size affects reproductive development and post– reproductive survival

In males, experimental body size manipulation impacted the timing of sexual development (LMM, LR = 14.19, p < 0.001; Fig. 2. c). Enlarged polyps started producing gonads significantly earlier than reduced ones (LMM, beta = –1.79, SE = 0.45, p < 0.001), but they did not differ from the control group (LMM, enlarged vs. control: beta = –0.38, SE = 1.18, p = 0.742). Furthermore, control and reduced male polyps did not differ from each other in their sexual development speed (LMM, control vs. reduced: beta = –1.41, SE = 1.18, p = 0.444). The number of testes was also impacted (Poisson GLMM, χ^2^ = 61.05, p < 0.001; Fig. 2. e). Enlarged male polyps produced a significantly higher number of testes than both control and reduced polyps (enlarged vs. control: beta = 0.24, SE = 0.06, p < 0.001; enlarged vs. reduced: beta = 0.62, SE = 0.08, p < 0.001). Furthermore, the number of testes in the reduced group was also significantly higher than in controls (Poisson GLMM, control vs. reduced: beta = 0.38, SE = 0.08, p < 0.001).

In females, experimental treatment affected the length of time required to produce the first egg (LMM, LR = 60.42, p < 0.001; Fig. 2. d). Enlarged female polyps took less time to develop eggs than reduced and control ones (LMM, enlarged vs. reduced: beta = –5.72, SE = 0.55, p < 0.001; enlarged vs. control: beta = –1.58, SE = 0.79, p = 0.039), and reduced polyps took more time to develop eggs than controls (LMM, control vs. reduced: beta = –4.13, SE = 0.77, p < 0.001). The number of eggs produced by females differed significantly between groups (Poisson GLMM, χ^2^ = 9.44, p = 0.009; Fig. 2. f). Enlarged female polyps had significantly more eggs than reduced ones (Poisson GLMM, enlarged vs. reduced: beta = 0.37, SE = 0.12, p = 0.007), but neither enlarged, nor reduced females differed from controls in egg number (Poisson GLMM, control vs. reduced: beta = 0.17, SE = 0.12, p = 0.166, enlarged vs. control: beta = 0.21, SE = 0.11, p = 0.132).

To see whether enlarged individuals have an increased reproductive allocation relative to their body size we analyzed the effect of experimental size manipulation on size–standardized relative gonad numbers. Experimental treatment impacted relative gonad numbers in both sexes (LMM, males, LR = 36.48, p < 0.001; females, LR = 15.85, p < 0.001). However, investigation of pairwise differences revealed that larger individuals had smaller size–specific gonad numbers. With one exception (female enlarged vs. control), these pairwise differences were statistically significant (LMM, males, control vs. reduced: beta = –0.30, SE = 0.06, p < 0.001; enlarged vs. control: beta = –0.16, SE = 0.06, p = 0.011; enlarged vs. reduced: beta = –0.46, SE = 0.07, p < 0.001; females, control vs. reduced: beta = –0.50, SE = 0.14, p < 0.001; enlarged vs. control: beta = –0.15, SE = 0.15, p = 0.293; enlarged vs. reduced: beta = –0.65, SE = 0.17, p < 0.001).

The survival rates were significantly impacted by body size manipulation in both of the male and female strain (binomial GLMM, male strain, χ^2^ = 13.58, p = 0.001; females, χ^2^ = 10.17, p = 0.006; Fig. 2. g, h). In the male strain, reduced polyps were more likely to survive compared to control or enlarged polyps (binomial GLMM, male strain, control vs. reduced: beta = –2.01, SE = 0.80, p = 0.024; enlarged vs. reduced: beta = –2.56, SE = 0.84, p = 0.007), but the difference between controls and enlarged polyps was not significant (binomial GLMM, male strain, enlarged vs. control: beta = –0.53, SE = 0.53, p = 0.315). In the female strain, reduced polyps had significantly higher survival rate than controls (binomial GLMM, control vs. reduced: beta = –1.70, SE = 0.56, p = 0.008) and the difference between reduced and enlarged polyps was marginally significant (binomial GLMM, enlarged vs. reduced: beta = –1.57, SE = 0.70, p = 0.052). Control and enlarged female polyps, on the other hand, did not differ from each other in survival (binomial GLMM, enlarged vs. control: beta = 0.13, SE = 0.67, 0.846).

## DISCUSSION

Using experimental manipulation, we here showed that body size *per se* has a causal effect in determining fecundity and longevity in a freshwater cnidarian. Particularly, a reduced body size was associated with delayed reproduction, lower reproductive output and improved survival rate in a male and female strain of *Hydra oligactis*. The experimental effects of body size manipulation aligned remarkably well with allometric patterns observed in a common garden experiment of 4 replicate samplings from a natural population over two years, suggesting that body size variation in itself is a major determinant of the natural diversity in sexual development and post–reproductive senescence in this species.

The higher fecundity of larger animals is a universal, well–documented relationship and one of the pillars of life history theory (Roff 1993; Stearns 2002; Shingleton 2011). Conversely, the relationship between body size and longevity is more complex and much less well understood. On the one hand, a large body size implies more resources that can be invested in fuelling physiological repair, potentially contributing to increased longevity. However, this was clearly not the case in our study as larger individuals were less likely to survive. On the other hand, a larger body is more costly to build and maintain, reducing resources available for physiological repair. That growth has viability costs is well known (Blanckenhorn 2000, Metcalf & Monaghan 2003) and these viability costs were hypothesized to explain the negative correlation between body size and longevity observed in several domesticated species (Rollo 2002, Austad 2010). Our data, however, suggest that a small size itself can lead to improved survival in a growth– independent manner, because the size differences in our experiment were achieved without altering growth rate. We can think of two different (though mutually not exclusive) explanations behind this phenomenon, as discussed below.

First, a larger body size implies a larger mass of tissue with proportionally more energetic requirements, all else being equal. While the animals enlarged in our experiment had a larger digestive cavity, their head region and tentacles (which contain most stinging cells and are critical for capturing food) were left intact during the grafting procedure, therefore it is unlikely that their food capture rate was increased. As a result, enlarged animals had to divide their ingested food among a larger tissue mass, reducing the amount of resources left for physiological maintenance and repair, which ultimately could have led to their reduced survival. This hypothesis predicts a negative relationship between total metabolic expenditure and physiological repair functions driven by body size variation. However, while the relationship between body size, metabolic rate and longevity is well described interspecifically (Speakman 2005, Killen et al. 2015), much less is known about intraspecific co–variation among these traits and the integration of physiological repair mechanisms into this co–variation.

Secondly, differences in post–reproductive survival of large and small individuals could have been the consequence of size–dependent reproductive decisions. For instance, if large individuals invest proportionally more of their resources into reproduction, that could drain resources available for survival. Such a hyperallometric investment in reproduction has been recently demonstrated in fish (Barneche et al. 2018) and might be common in the natural world (Marshall & White 2019). This hypothesis would predict that relative (size–standardized) gonad number in hydra increases with body size. Contrarily, we found a negative relationship between relative gonad number and body size for both males and females in the common garden animals and the size–manipulated polyps. Hence, reproductive investment in *H. oligactis* appears to be hypoallometric, instead of hyperallometric and size–dependent reproductive investment is unlikely to explain the increased survival rate of smaller individuals.

On the other hand, we did observe size-dependent adjustment of sexual development speed. In both the common garden animals and size–manipulated polyps, smaller individuals took more time to start gonadogenesis than larger individuals. Since in the laboratory animals are kept on constant food, this could have enabled small individuals to accumulate more resources, ultimately enabling them to attain increased survival. Why do small individuals but not large ones delay reproduction? In the case of males, this could be explained by intrasexual competition: in their natural habitat, *H. oligactis* males appear earlier in the season (Sebestyén et al. 2018) and likely compete to be first to fertilize females. A small male is expected to lose out in competition for fertilization, due to its lower number of reproductive organs. Hence small individuals might receive a greater fitness payoff by investing into physiological repair to increase the chances of overwinter survival and participation in a second reproductive season. In females, the situation is much less clear, because we do not expect intrasexual competition in their case. However, delayed sexual development and increased investment into survival in small females could be caused, for instance, by sexually antagonistic selection (selection on males causing correlated changes in females; Bonduriansky et al. 2008). This hypothesis is all the more likely since sex– change is known to occur in this species (Miklós et al. 2019 and this study), therefore the same genotype might have experienced selection both as a female and as a male. Alternatively, females might be time-constrained as well (e.g. to produce eggs before the water freezes, or to match egg maturation to male availability), but if small females are less able to produce high– quality eggs due to their reduced condition, it might be optimal for them to invest in survival instead.

While multiple mechanisms might be involved, our data clearly show that body size itself influences reproductive decisions, potentially explaining the differential survival rate of small and large individuals. Since the relationship between body size and reproductive decisions is well established in the animal world (Reiss 1987) and reproductive decisions have carry–over effects on survival in animals ranging from nematodes and fruit flies to humans (Hsin & Kenyon 1999, Flatt 2011, Min et al. 2012), the link between body size, reproduction and senescence might be a common one in the natural world.

What could be the physiological mechanism through which body size influences other life history traits in hydra? According to a recent study, hydra polyps assess and control their body size through the Wnt and TGF–β signaling pathways, which regulate cell proliferation and differentiation in the body column (Mortzfeld et al. 2019). Wnt signaling is one of the major effectors of the head organizer, a population of cells in the head region which emits diffusible signals that regulate body plan formation and budding of hydra polyps (Hobmeyer et al. 2000; Shimizu 2012). Experimental manipulation of body size is likely to alter this system by changing the diffusion distance and effectivity of small regulatory molecules emitted from the head region, which could alter the differentiation of gametes and hence reproductive investment. Because gametes in hydra derive from the same stock of interstitial stem cell populations that also give rise to somatic derivatives (such as the stinging cells necessary for food capture; Nishimiya– Fujisawa & Kobayashi 2018), any shift in the differentiation of stem cells to gametes is likely to reduce cell populations involved in somatic maintenance, thereby impacting survival (Sebestyén et al. 2018).

Alternatively, interaction between germ cell precursors is another mechanism that could influence timing and intensity of gamete maturation. In a number of organisms reproductive investment and somatic maintenance are regulated by germline signals that increase reproduction at the expense of survival (e.g. Hsin & Kenyon 1999; Flatt et al. 2008). Interactions between germ cell precursors are known to exist in the germline of hydra polyps as well (Miller et al. 2000; Nishimiya–Fujisawa & Kobayashi 2018). In our experiment, an increased size means more germ cells and potentially more germline signals which could shift the balance of differentiation from somatic to sexual investment. We note however, that the reduced size–specific gonad number of larger individuals speaks against this possibility. The exact physiological mechanisms behind size–specific survival in hydra remain to be addressed in the future but they provide exciting opportunities for deciphering the physiology of size–dependent senescence.

In conclusion, by exploiting the ability of hydra polyps to incorporate tissue from other polyps, we were able to experimentally alter body size in *H. oligactis*, a freshwater cnidarian with experimentally inducible sex and post–reproductive senescence. Size manipulation altered the timing of sexual development and fecundity, and impacted post–reproductive survival, such that smaller polyps had improved survival. Our data provide experimental support for fundamental hypotheses in life history evolution that were based on a solid theoretical background of allometric relationships and decades of correlative data collection, but lacked an experimental approach. The *Hydra* model system might be a fruitful model system to further understand allometric patterns in life history traits.

## ACKNOWLEDGEMENTS

This study was supported by NKFIH grant FK 124164. BA was supported by the Hungarian State and the European Union under the EFOP–3.6.1 project. ZB was financed by the Higher Education Institutional Excellence Programme (NKFIH–1150–6/2019) of the Ministry of Innovation and Technology in Hungary, within the framework of the DE–FIKP Behavioral Ecology Research Group thematic program of the University of Debrecen. JT was supported by a János Bolyai Research Scholarship of the Hungarian Academy of Sciences and the ÚNKP New National Excellence Program of the Hungarian Ministry of Innovation and Technology (ÚNKP–19–4). We thank Máté Miklós, Flóra Sebestyén, Beatrix Kozma, Dávid Tenkei, Réka Gergely and Erzsébet Ágnes Nehéz for lab assistance and Erzsébet Ágnes Nehéz for producing photographs on Fig. 3.

## REFERENCES

Andersson, M. (1994). Sexual selection. Princeton University Press. Princeton, New Jersey.

Austad S.N. (2010) Animal size, metabolic rate, and survival, among and within species. In: Wolf N. (ed.) The Comparative biology of aging. pp. 27–43. Springer, Dordrecht

Allaine, D., Pontier, D., Gaillard, J. M., Lebreton, J. D., Trouvilliez, J., & Clobert, J. (1987). The relationship between fecundity and adult body weight in homeotherms. Oecologia, 73(3), 478–480.

Arak, A. (1988). Sexual dimorphism in body size: a model and a test. Evolution, 42(4), 820–825.

Barneche, D. R. and Robertson, D. R. and White, C. R. and Marshall, D. J. (2018). Fish reproductive–energy output increases disproportionately with body size. Science, 360(6389), 642–645.

Bartke, A. (2017). Somatic growth, aging, and longevity. NPJ aging and mechanisms of disease, 3(1), 1–6.

Bates, D., Maechler, M., Bolker, B., Walker, S. (2015). Fitting Linear Mixed–Effects Models Using lme4. Journal of Statistical Software, 67(1), 1–48.

Blanckenhorn, W. U. (2000). The evolution of body size: what keeps organisms small?. The quarterly review of biology, 75(4), 385–407.

Bonduriansky, R., Maklakov, A., Zajitschek, F., & Brooks, R. (2008). Sexual selection, sexual conflict and the evolution of ageing and life span. Functional ecology, 22(3), 443–453.

Bosch, T. C. (2007). Why polyps regenerate and we don’t: towards a cellular and molecular framework for Hydra regeneration. Developmental biology, 303(2), 421–433.

Darwin, C. R. (1874). The descent of man, and selection in relation to sex. 2d ed. Appleton, New York.

Damuth, J. (2007). A macroevolutionary explanation for energy equivalence in the scaling of body size and population density. The American Naturalist 169(5), 621–631.

Flatt, T., Min, K. J., D’Alterio, C., Villa–Cuesta, E., Cumbers, J., Lehmann, R., Jones, D. L. & Tatar, M. (2008). Drosophila germ–line modulation of insulin signaling and lifespan. Proceedings of the National Academy of Sciences, 105(17), 6368–6373.

Flatt, T. (2011). Survival costs of reproduction in Drosophila. Experimental gerontology, 46(5), 369–375.

Forsman, A. (1993). Survival in relation to body size and growth rate in the adder, Vipera berus. Journal of Animal Ecology, 647–655.

Gaillard, J. M., Festa–Bianchet, M., Delorme, D., & Jorgenson, J. (2000). Body mass and individual fitness in female ungulates: bigger is not always better. Proceedings of the Royal Society of London. Series B: Biological Sciences, 267(1442), 471–477.

He, Q., Morris, B. J., Grove, J. S., Petrovitch, H., Ross, W., Masaki, K. H., Rodriguez, B., Chen, R., Donlon, T. A., Willcox, D. C. & Willcox, B. J. (2014). Shorter men live longer: association of height with longevity and FOXO3 genotype in American men of Japanese ancestry. PLoS One, 9(5), e94385.

Hendriks, A. J. & Mulder, C. (2008). Scaling of offspring number and mass to plant and animal size: model and meta–analysis. Oecologia 155, 705–716.

Hobmayer, B., Rentzsch, F., Kuhn, K., Happel, C. M., von Laue, C. C., Snyder, P., Rothbächer, U. & Holstein, T. W. (2000). WNT signalling molecules act in axis formation in the diploblastic metazoan Hydra. Nature, 407(6801), 186–189.

Hothorn, T., Bretz, F. & Westfall, P. (2008). Simultaneous inference in general parametric models. Biometrical Journal, 50(3), 346–363.

Killen, S. S., Atkinson, D., & Glazier, D. S. (2010). The intraspecific scaling of metabolic rate with body mass in fishes depends on lifestyle and temperature. Ecology letters, 13(2), 184–193.

Kirkwood, T. B., & Rose, M. R. (1991). Evolution of senescence: late survival sacrificed for reproduction. Philosophical Transactions of the Royal Society of London. Series B: Biological Sciences, 332(1262), 15–24.

Kraus, C., Pavard, S., & Promislow, D. E. (2013). The size–life span trade–off decomposed: why large dogs die young. The American Naturalist, 181(4), 492–505.

Marshall, D. J., & White, C. R. (2019). Have we outgrown the existing models of growth?. Trends in ecology & evolution, 34(2), 102–111.

Martínez, D. E., (1998). Mortality patterns suggest lack of senescence in hydra. Experimental Gerontology, 33, 217–225.

McCulloch, D., & Gems, D. (2003). Body size, insulin/IGF signaling and aging in the nematode *Caenorhabditis elegans*. Experimental gerontology, 38(1–2), 129–136.

Metcalfe, N. B., & Monaghan, P. (2003). Growth versus lifespan: perspectives from evolutionary ecology. Experimental gerontology, 38(9), 935–940.

Miller, M. A., Technau, U., Smith, K. M., & Steele, R. E. (2000). Oocyte development in Hydra involves selection from competent precursor cells. Developmental biology, 224(2), 326–338.

Min, K. J., Lee, C. K., & Park, H. N. (2012). The lifespan of Korean eunuchs. Current Biology, 22(18), R792–R793.

Miklós, M., Laczkó, L., Sramkó, G., Sebestyén, F., Barta, Z. & Tökölyi, J. (2019). Population genomic structure underlying extremely divergent life history strategies in hydra. Preprint. https://www.doi.org/10.13140/RG.2.2.34448.87044.

Mortzfeld, B. M., Taubenheim, J., Klimovich, A. V., Fraune, S., Rosenstiel, P., & Bosch, T. C. (2019). Temperature and insulin signaling regulate body size in Hydra by the Wnt and TGF–beta pathways. Nature communications, 10(1), 1–13.

Nishimiya–Fujisawa, C., & Kobayashi, S. (2018). Roles of germline stem cells and somatic multipotent stem cells in Hydra sexual reproduction. In: Kobayashi, K., Kitano, T., Iwao, Y. & Kondo, M. (eds.) Reproductive and developmental strategies (pp. 123–155). Springer, Tokyo.

Peters, R. H. (1983). The ecological implications of body size. Cambridge University Press. Cambridge, UK.

Pinheiro, J., Bates, D., DebRoy, S., Sarkar, D., R Core Team (2020). _nlme: Linear and Nonlinear Mixed Effects Models_. R package version 3.1–144, <URL: https://CRAN.R-project.org/package=nlme>.

Promislow, D. E., & Harvey, P. H. (1990). Living fast and dying young: A comparative analysis of life-history variation among mammals. Journal of Zoology, 220(3), 417–437.

R Core Team (2020). R: A language and environment for statistical computing. R Foundation for Statistical Computing, Vienna, Austria. URL https://www.R-project.org/.

Reiss, M. J. (1987). The intraspecific relationship of parental investment to female body weight. Functional Ecology, 105–107.

Roff, D. (1992). Evolution of life histories: theory and analysis. Chapman & Hall, New York, NY.

Rollo, C. D. (2002). Growth negatively impacts the life span of mammals. Evolution & development, 4(1), 55–61.

Salminen, A., & Kaarniranta, K. (2010). Insulin/IGF–1 paradox of aging: regulation via AKT/IKK/NF–κB signaling. Cellular signalling, 22(4), 573–577.

Samaras, T. T., Elrick, H., & Storms, L. H. (2003). Is height related to longevity?. Life sciences, 72(16), 1781–1802.

Schaible, R., Scheuerlein, A., Dańko, M. J., Gampe, J., Martínez, D. E., & Vaupel, J. W. (2015). Constant mortality and fertility over age in Hydra. Proceedings of the National Academy of Sciences, 112(51), 15701–15706.

Schneider, C. A., Rasband, W. S., & Eliceiri, K. W. (2012). NIH Image to ImageJ: 25 years of image analysis. Nature methods, 9(7), 671–675.

Sebestyén, F., Barta, Z., & Tökölyi, J. (2018). Reproductive mode, stem cells and regeneration in a freshwater cnidarian with postreproductive senescence. Functional Ecology, 32(11), 2497–2508.

Sebestyén, F., Miklós, M., Iván, K., & Tökölyi, J. (*in press*). Age–dependent plasticity in reproductive investment, regeneration capacity and survival in a partially clonal animal (*Hydra oligactis*). Journal of Animal Ecology.

Shimizu, H. (2012). Transplantation analysis of developmental mechanisms in Hydra. International Journal of Developmental Biology, 56(6–7–8), 463–472.

Shingleton, A. W. (2011). Evolution and the regulation of growth and body size. In: Flatt, T. & Heyland, A. (eds.) Mechanisms of life history evolution, pp 43–55. Oxford University Press, Oxford.

Speakman, J. R. (2005). Body size, energy metabolism and lifespan. Journal of Experimental Biology, 208(9), 1717–1730.

Stearns, S. C. (1992). The evolution of life histories. Oxford University Press, Oxford.

Székely, T., Reynolds, J. D., & Figuerola, J. (2000). Sexual size dimorphism in shorebirds, gulls, and alcids: the influence of sexual and natural selection. Evolution, 54(4), 1404–1413.

Tökölyi, J., Ősz, Z., Sebestyén, F., & Barta, Z. (2017). Resource allocation and post– reproductive degeneration in the freshwater cnidarian Hydra oligactis (Pallas, 1766). Zoology, 120, 110–116.

Tökölyi, J., Gergely, R., & Miklós, M. (2020). Seasonal variation in sexual readiness in a facultatively sexual freshwater cnidarian with diapausing eggs. Preprint. https://doi.org/10.1101/2020.05.27.119123

Tomczyk, S., Suknovic, N., Schenkelaars, Q., Wenger, Y., Ekundayo, K., Buzgariu, W., Bauer, Ch., Fisher, K., Austad, S. & Galliot, B. (2020). Deficient autophagy in epithelial stem cells drives aging in the freshwater cnidarian Hydra. Development, 147:dev177840.

Yoshida, K., Fujisawa, T., Hwang, J. S., Ikeo, K., & Gojobori, T. (2006). Degeneration after sexual differentiation in hydra and its relevance to the evolution of aging. Gene, 385, 64–70.

